# Outer mitochondrial membrane E3 Ub ligase MARCH5 controls mitochondrial steps in peroxisome biogenesis

**DOI:** 10.1101/2023.08.31.555756

**Authors:** Nicolas Verhoeven, Yumiko Oshima, Etienne Cartier, Albert Neutzner, Liron Boyman, Mariusz Karbowski

## Abstract

Peroxisome *de novo* biogenesis requires yet unidentified mitochondrial proteins. We report that the outer mitochondrial membrane (OMM)-associated E3 Ub ligase MARCH5 is vital for generating mitochondria-derived pre-peroxisomes. MARCH5 knockout results in accumulation of immature peroxisomes and lower expression of various peroxisomal proteins. Upon fatty acid-induced peroxisomal biogenesis, MARCH5 redistributes to newly formed peroxisomes; the peroxisomal biogenesis under these conditions is inhibited in MARCH5 knockout cells. MARCH5 activity-deficient mutants are stalled on peroxisomes and induce accumulation of peroxisomes containing high levels of the OMM protein Tom20 (mitochondria-derived pre-peroxisomes). Furthermore, depletion of peroxisome biogenesis factor Pex14 leads to the formation of MARCH5- and Tom20-positive peroxisomes, while no peroxisomes are detected in Pex14/MARCH5 dko cells. Reexpression of WT, but not MARCH5 mutants, restores Tom20-positive pre-peroxisomes in Pex14/MARCH5 dko cells. Thus, MARCH5 acts upstream of Pex14 in mitochondrial steps of peroxisome biogenesis. Our data validate the hybrid, mitochondria-dependent model of peroxisome biogenesis and reveal that MARCH5 is an essential mitochondrial protein in this process.

**Summary:** The authors found that mitochondrial E3 Ub ligase MARCH5 controls the formation of mitochondria-derived pre-peroxisomes. The data support the hybrid, mitochondria-dependent model of peroxisome biogenesis and reveal that MARCH5 is an essential mitochondrial protein in this process.

## Introduction

Extensive evidence supports the vital role of the ubiquitin (Ub) proteasome system (UPS) in mitochondrial function (Benard et al., 2010; Chen et al., 2013; Cherok et al., 2017; Karbowski et al., 2007; Karbowski and Youle, 2011; Neutzner et al., 2012; Oshima et al., 2021; Tanaka et al., 2010; Xu et al., 2016; Xu et al., 2011). In addition to the well-established elimination of dysfunctional mitochondria through E3 Ub ligase Parkin-dependent mitochondria-specific autophagy (mitophagy), the UPS also controls the homeostasis of healthy mitochondria. MARCH5 (Mitol) is an integral E3 Ub ligase of the outer mitochondrial membrane (OMM) implicated in many mitochondrial pathways, including regulation of mitochondrial fission and fusion, mitochondrial steps in apoptosis, and removal of disease-linked toxic proteins from the mitochondria (Chen et al., 2017; Cherok et al., 2017; Hussain et al., 2021; Karbowski et al., 2007; Koyano et al., 2019; Park et al., 2010; Park et al., 2014; Shiiba et al., 2021; Sugiura et al., 2013; Sugiura et al., 2011; Xu et al., 2016; Yoo et al., 2019). MARCH5 expression is regulated by peroxisome proliferator-activated receptor-γ (PPAR γ) and negatively correlates with fat mass across a panel of genetically diverse mouse strains, including a common obesity model (*ob/ob*) in mice (Bond et al., 2019). This relationship has also borne out in people suffering from obesity and a large cohort with metabolic syndrome (Bond et al., 2019), linking MARCH5 with regulating energy metabolism. Consistent with the metabolic function of MARCH5, the Genome-wide associated loci prioritization (Open Targets Genetics) shows a high-confidence prediction of MARCH5 mutations as a likely causal gene for obesity and type II diabetes. However, the role and mechanism of MARCH5 in cellular energy metabolism are poorly understood. Zheng et al. (Zheng et al., 2022) showed that, in addition to the OMM, a subset of MARCH5 also localizes to the peroxisomes. Peroxisomes are abundant, dynamic single-membrane organelles that host diverse biochemical pathways such as ether-phospholipid biosynthesis, bile acid synthesis, glyoxylate metabolism, and amino acid catabolism. Peroxisomes also play a central role in fatty acid β-oxidation (Fujiki et al., 2020; Islinger et al., 2018) with an apparent preference for chain-shortening of very long-chain fatty acids, but can also metabolize medium- and long-chain fatty acids such as palmitate (Violante et al., 2019) that are typically oxidized by mitochondria, thereby providing energy intermediate donors for mitochondrial ATP generation (Houten et al., 2020). While the function of MARCH5 in peroxisome-specific autophagy (pexophagy) was also proposed (Zheng et al., 2022), the scope and mechanism of MARCH5, and potential crosstalk in the control of mitochondrial and peroxisomal pathways are not known.

The functional and molecular connection between mitochondria and peroxisomes is further underscored by the fact that many proteins initially thought to exclusively localize to the mitochondria were later found on peroxisomes, including fission factors dynamin-related protein 1 (Drp1), Fis1 and mitochondrial fission factor (Mff), AAA-ATPase ATAD1 (Msp1), MARCH5 (Castanzo et al., 2020; Gandre-Babbe and van der Bliek, 2008; Koch et al., 2005; Zheng et al., 2022). On the other hand, several peroxisomal proteins, such as Pex26, Pex14, and Pex3, localize to the OMM, especially when peroxisomal biogenesis machinery is dysfunctional (Castanzo et al., 2020; Chen et al., 2014; Nuebel et al., 2021; Okreglak and Walter, 2014). The accumulation of peroxisomal proteins on the OMM, which was broadly considered a mis-localization, could reflect the recently discovered mitochondrial role in peroxisomal biogenesis (Schrader and Pellegrini, 2017; Sugiura et al., 2017). New peroxisomes in the cell can form by the growth and division of pre-existing organelles but can also emerge *de novo.* In the *de novo* biogenesis pathway, a subset of pre-peroxisomes develops from the mitochondria, and some peroxisomal biogenesis factors, including Pex3 and Pex14, could have a specific mitochondrial function in this process (Sugiura et al., 2017). The mature, metabolically active peroxisomes are then formed by the fusion of ER- and mitochondria-derived pre-peroxisomes (hybrid model of peroxisome biogenesis) (Schrader and Pellegrini, 2017; Sugiura et al., 2017). This line of thought is further substantiated in peroxisome biogenesis disorders (PBD), such as Zellweger’s syndrome, as patient-derived cells accumulate peroxisome proteins on the OMM (Nuebel et al., 2021). The mitochondrial proteins required for mitochondrial steps in peroxisome biogenesis are yet unknown. In addition to the coordinated biogenesis and fatty acid oxidation, mitochondria and peroxisomes also interplay in other pathways, such as the control of redox homeostasis, anti-viral signaling, and apoptosis (Fransen et al., 2017; Tanaka et al., 2019). The integration of pathways coregulated by mitochondria and peroxisomes could partly be achieved through mitochondria-peroxisome contact sites, as reported in Leydig cells’ steroid biosynthesis (Fan et al., 2016).

Here, we report the data supporting the central role of MARCH5 in peroxisome biogenesis and, thereby, cellular adaptation to distinct availability of energy substrates. Our data show that MARCH5 acts upstream of Pex14 in formation of mitochondria-derived pre-peroxisomes.

## Results and Discussion

### Activity-dependent peroxisomal localization of MARCH5

Using superresolution Airyscan imaging we tested subcellular localization of MYC-tagged WT MARCH5 (MYC-WT MARCH5), RING domain activity-deficient MARCH5 (MYC-MARCH5^H43W^; (Cherok et al., 2017; Karbowski et al., 2007)) and the C-terminal truncation mutant (MYC-MARCH5^ΔCT^; (Cherok et al., 2017)) in MARCH5 knockout (MARCH5^−/−^) HeLa cells. MARCH5^H43W^ and MARCH5^ΔCT^ mutants were chosen because of their reported inhibitory effects on MARCH5 activity (Cherok et al., 2017; Karbowski et al., 2007; Xu et al., 2016). Cells were transfected with respective constructs followed by immunostaining to detect peroxisomes (Pex14), MARCH5 (anti-MYC), and the outer mitochondrial membrane (OMM; Tom20). The data show that while MYC-WT MARCH5 localized mainly to the mitochondria, with occasional peroxisomal colocalization (Fig. 1A,D), MYC-MARCH5^H43W^ and MYC-MARCH5^ΔC^ were invariably associated with peroxisomes (Fig. 1D). Since these mutants show reduced ubiquitination activity toward MARCH5 substrates (Cherok et al., 2017; Xu et al., 2016), it is likely that inhibition of MARCH5 E3 Ub ligase activity increases association of MARCH5 with peroxisomes. Furthermore, in contrast to MYC-WT MARCH5 expressing cells that show negligible levels of the outer mitochondrial membrane marker Tom20 colocalizing with Pex14 (Fig. 1A, D), cells expressing MARCH5 mutants showed high levels of Tom20 colocalization with Pex14 (Fig. 1B-D). These data suggest that MARCH5 activity is implicated in the biogenesis of the mitochondria-derived pre-peroxisomes. It is likely that MARCH5 E3 Ub ligase activity mediates early steps in mitochondrial pre-peroxisome formation and its inhibition induces abnormal accumulation of the OMM protein Tom20-containing immature peroxisomes, associated with retention of inactive MARCH5 on Pex14-positive peroxisomes. Consistent with this notion, the formation of mitochondria derived pre-peroxisomes was also proposed to be regulated by Pex14 (Schrader and Pellegrini, 2017; Sugiura et al., 2017).

**Figure 1:**
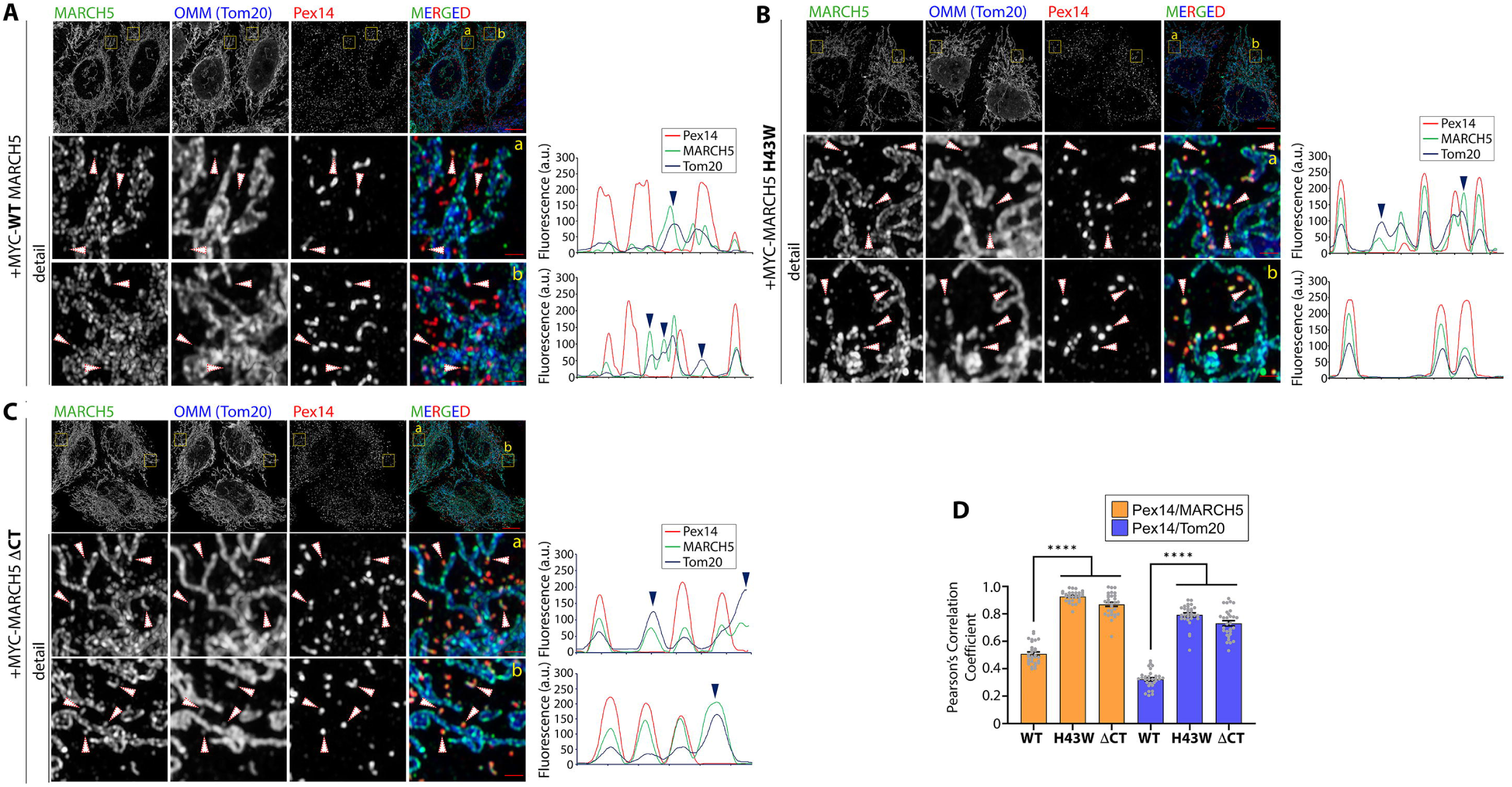
Peroxisomal localization of MARCH5. Typical Airyscan superresolution images of MARCH5^−/−^ HeLa cells transfected with MYC-WT MARCH5 (**A**), MYC-MARCH5 ^H43W^ (**B**), or MARCH5^ΔCT^ (**C**) immunostained to detect peroxisomes (Pex14; red on merged images), MARCH5 (anti-MYC Ab; green on merged images), and outer mitochondrial membrane (Tom20; blue on merged images). Detail images are from the areas marked with yellow rectangles. White arrowheads indicate typical Pex14, and corresponding MARCH5 and Tom20 staining patterns. Fluorescence profiles are shown on the right sides of the images. Blue arrowheads indicate examples of mitochondrial Tom20 and MARCH5 fluorescence. Bars: 10μm and 1μm in detail images. (**D**) Pearson’s colocalization coefficients representing colocalization of peroxisomes (Pex14) with MARCH5 (orange), or peroxisomes (Pex14) with Tom20 (blue). Mean+/−SEM. N=30 cells *per* condition. One-way ANOVA. **** p<0.0001.

### MARCH5 deficiency affects peroxisome homeostasis

The recent work by Zheng *et al*. (Zheng et al., 2022), also shows peroxisomal localization of WT MARCH5, and proposed the role of MARCH5 in peroxisome-specific autophagy (pexophagy). However, a detailed analysis of peroxisome status in MARCH5 deficient cells has yet to be reported. We used the quantitative image analysis to determine the effect of MARCH5 knockout on peroxisome number, size, and maturation. The data show complete colocalization of peroxisome biogenesis factor PMP70 with a marker of “mature” peroxisomes – Catalase – in WT cells (Fig. 2A). On the other hand, in MARCH5^−/−^ cells, many PMP70-positive peroxisomes were deficient of Catalase staining (Fig. 2B; blue arrowheads). Quantifying peroxisome number also indicates reduced Catalase-positive peroxisomes, compared to PMP70- or Pex14-positive peroxisomes, in both MARCH5 deficient HeLa and HCT116 cells. Specifically, there are ~20% and 22% fewer Catalase-positive peroxisomes *per* cell compared to the number of PMP70-peroxisomes in MARCH5^−/−^ HeLa and HCT116 cells, respectively (Fig. 2C). On the other hand, no difference between PMP70- and Catalase-positive peroxisomes was apparent in WT HeLa and HCT116 cells (Fig. 2C). Furthermore, peroxisomes in MARCH5^−/−^ cells are smaller than in WT cells (the average size of PMP70-positive peroxisomes is 0.089+/−0.013μm^2^ or 0.094+/−0.015μm^2^ *versus* 0.067+/−0.012μm^2^ or 0.076+/−0.011μm^2^ in WT HeLa and WT HCT116 and MARCH5-deficient cells, respectively; Fig. 2D). Since similar data were found in two independent MARCH5^−/−^ cell models, one can conclude that MARCH5 controls peroxisome size and number. The fact that many peroxisomes in MARCH5^−/−^ did not contain detectable levels of Catalase (Fig. 2B) also supports the possibility that peroxisome biogenesis stalled before the Catalase import. Consistent with this notion, cell fractionation shows accumulation of Catalase in cytosolic fractions from MARCH5^−/−^ cells, compared to WT cells (Fig. 2E; red arrowheads).

**Figure 2:**
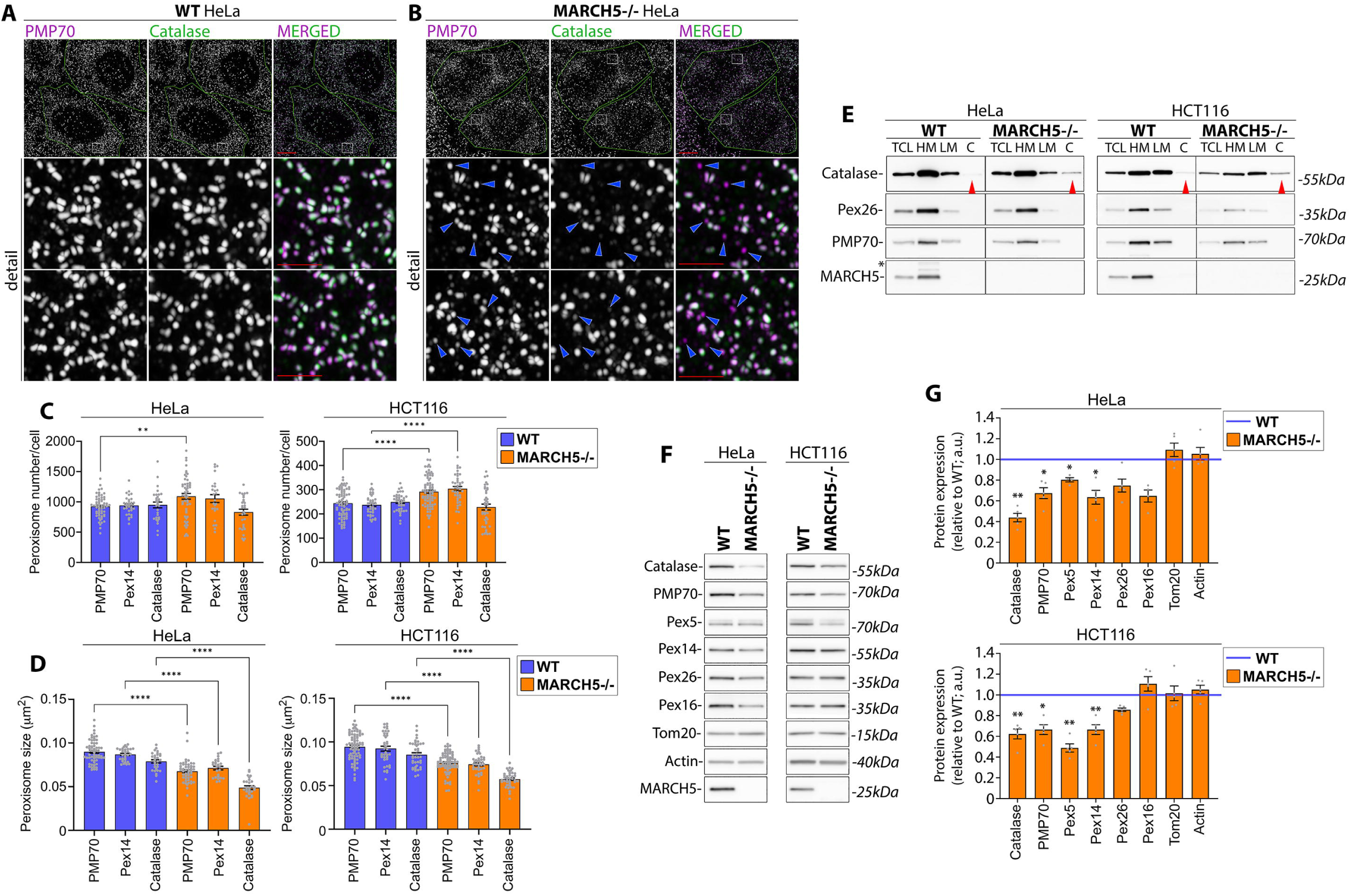
Defective peroxisomes in MARCH5^−/−^ cells. WT (**A**) and MARCH5^−/−^ (**B**) HeLa cells were immunostained to detect PMP70 (peroxisome biogenesis factor; purple) and Catalase (marker of metabolically proficient, “mature” peroxisomes; green). Arrowheads in detail images in **B** indicate PMP70-positive, Catalase-negative peroxisomes. Bars: 10μm and 2μm in detail images. (**C**) Number of Pex14-, PMP70- and Catalase-positive peroxisomes *per* cell in WT and MARCH5^−/−^ HeLa (left) and HCT116 (right) cells. Mean+/−SEM. N=62 (WT HeLa; PMP70), 52 (MARCH5^−/−^ HeLa: PMP70), 29 (WT HeLa, Catalase), 33 (WT HeLa; Pex14). Mean+/−SEM. One-way ANOVA. ** p<0.01, **** p<0.0001 (**D**) Size of Pex14-, PMP70- and Catalase-positive peroxisomes in WT and MARCH5^−/−^ HeLa (left) and HCt116 (right) cells. Mean+/−SEM. One-way ANOVA. **** p<0.0001. Data were pulled from 3 independent experiments. (**E**) WT and MARCH5^−/−^ HeLa (left panels) and HCT116 (right panels) cells were fractionated into mitochondria-enriched heavy membrane (HM), light membrane (LM) and cytosolic fractions (C) followed by Western blot to detect proteins indicated in the figure. TCL: total cell lysate. * Indicates a nonspecific signal remaining after membrane reblotting. Red arrowheads indicate Catalase levels in cytosolic fractions. (**F**) Total cell lysates from WT and MARCH5^−/−^ HeLa and HCT116 cells indicated in the figure were analyzed for expression levels of peroxisomal proteins. Actin and Tom20 are loading controls. (**G**) Densitometric evaluation of the expression of peroxisomal proteins treated as shown in **F**. Mean+/−SEM. N=3-5 independent experiments. One-Sample t-test. ** p<0.01, * p<0.05.

Previous reports show that the depletion of one peroxisome biogenesis factor leads to altered steady-state levels of other peroxisomal proteins (Ott et al., 2023; Wei et al., 2021). We tested the effect of MARCH5 deficiency on the expression of peroxisomal proteins. Total cell lysates from WT and MARCH5^−/−^ HeLa and HCT116 cells were analyzed by Western blot (Fig. 2F, G) to detect the representative peroxisomal proteins. The data show a consistent reduction in the expression of peroxisome biogenesis factors PMP70, Pex5, and Pex14, and the marker of mature peroxisomes Catalase in both HeLa and HCT116 MARCH5^−/−^ as compared to WT cells (Fig. 2F). On the other hand, expressions of Pex16 and Pex26 were only reduced in MARCH5^−/−^ HeLa cells. The densitometric quantification of several independent experiments is shown in Figure 2G. These data indicate that depletion of MARCH5 leads to reduced expression of peroxisomal proteins, resembling similar effects of the deficiencies in established peroxisome biogenesis factors (Ott et al., 2023; Wei et al., 2021). Together with the accumulation of Catalase-deficient peroxisomes (Fig. 2B) and reduced peroxisome size in MARCH5^−/−^ cells (Fig. 2D), these data suggest that MARCH5 is vital for peroxisome biogenesis.

### MARCH5 controls peroxisome biogenesis under fatty acid- and OXPHOS-dependent growth

The current understanding of the role and mechanisms by which the UPS factors, such as MARCH5, regulate mitochondrial and peroxisomal function is mostly based on studies of cells grown in standard 25mM glucose (high glucose) media. Under high glucose growth, cells use mitochondria as biosynthetic hubs but rely primarily on glycolysis to generate ATP. Therefore, under such conditions, mitochondria display relatively low bioenergetic activity (Ahn and Metallo, 2015). Even primary, normally OXPHOS-dependent cells adapt to glycolysis as a major source of ATP when high glucose levels are available. This is exemplified by rat hepatocytes, which, after a few days of culture in high glucose media, shift their ATP generation from OXPHOS to glycolysis (Fu et al., 2013). Switching cells to low or glucose-deficient growth conditions and providing OXPHOS-stimulating substrates is predicted to increase the utilization of mitochondria for ATP generation and activation of peroxisomal metabolic pathways, especially when fatty acids are used as OXPHOS substrates. We set up cell growth conditions in which cells predominantly rely on OXPHOS as the source of ATP generation. This cell culture model consists of a glucose-free medium, supplemented with 4.5mM galactose and 25μM palmitoyl-L-carnitine (pLcar), as a donor of OXPHOS substrates. In addition to glucose exclusion, pyruvate, a final metabolite in the glycolysis pathway, was removed from this cell culture medium (OXPHOS medium) to promote mitochondrial energy reliance on fatty acid oxidation. Since MARCH5 is a peroxisome proliferator-activated receptorγ (PPARγ) target gene and appears to regulate adipocyte metabolism (Bond et al., 2019), applying pLcar as an OXPHOS substrate donor is highly relevant. Additionally, in mammalian cells and in *S. cerevisiae*, exposure to fatty acids induces the expression of genes encoding various peroxisomal proteins, causing the organelles’ biogenesis and/or maturation.

WT HeLa cells shifted from high (25mM) glucose to the “OXPHOS medium” (Fig. 3A for experimental protocol) showed increased levels of OXPHOS proteins, including SDHB (respiratory complex II), COX-2 (complex IV), and UQCRC-2 (complex III) (Fig. 3B,C), and peroxisomal proteins Pex14 and Catalase (Fig. 3B,C). Thus, through increased expression or inhibited degradation of relevant proteins, mitochondrial and peroxisomal biogenesis is likely initiated by the shift into the OXPHOS medium. An increased number of OXPHOS complexes and/or peroxisomes may facilitate energy generation shift from glycolysis into fatty acid-supported OXPHOS. MARCH5^−/−^ cells show a similar to WT cells increase in expression of OXPHOS proteins (Fig. 3B,C), indicating that culture in the OXPHOS medium may induce mitochondrial biogenesis, which does not require MARCH5 activity. On the other hand, the increase in expression of peroxisomal markers, Catalase and Pex14, was significantly lower in MARCH5^−/−^ cells compared to WT HeLa cells (Fig. 3B,C). Quantification of Airyscan images also revealed an increase in peroxisome number in OXPHOS media-treated WT, but not MARCH5^−/−^ cells (Fig. 3D). Thus, in addition to controlling peroxisomal homeostasis under standard 25mM glucose-supported growth, MARCH5 appears to be required for OXPHOS medium-induced peroxisome biogenesis.

**Figure 3.**
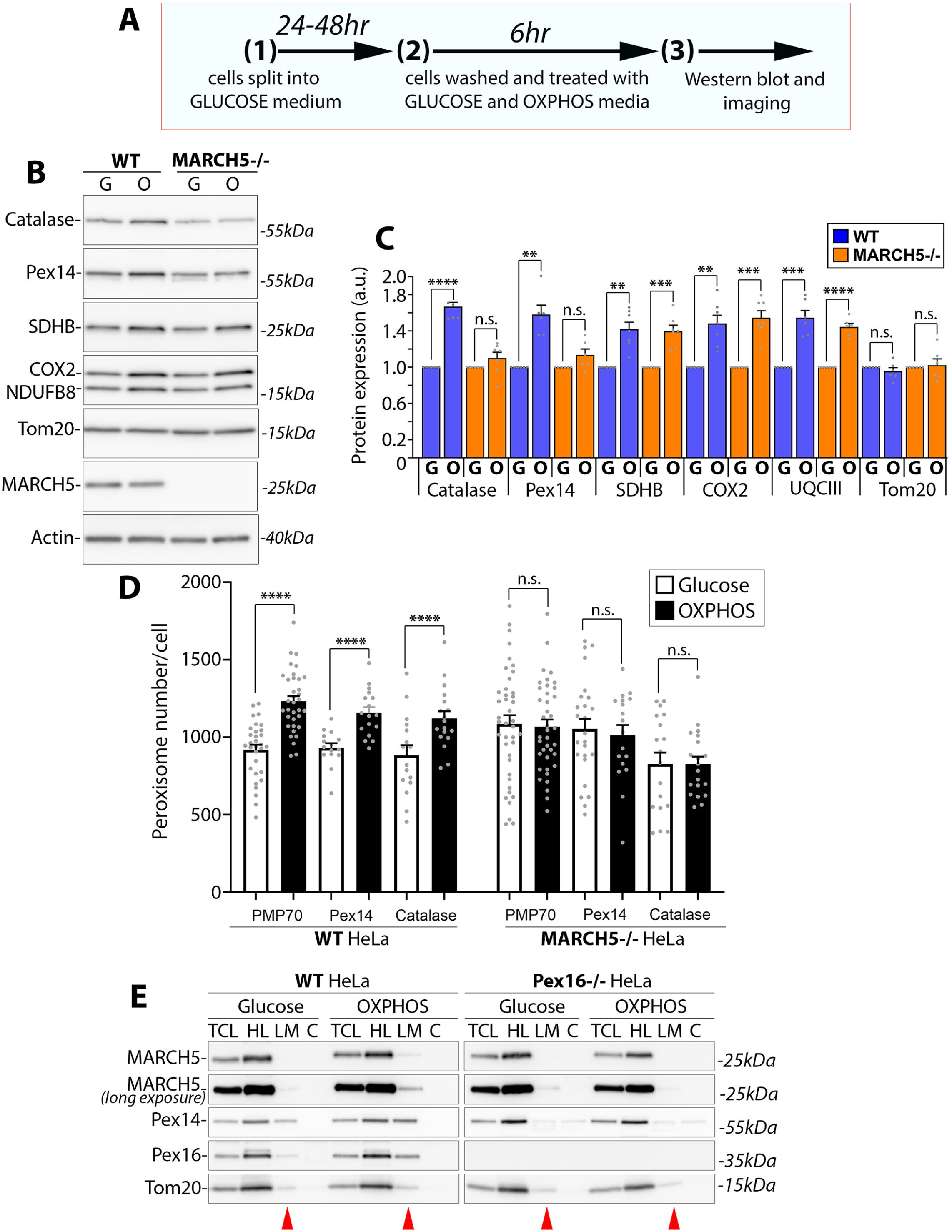
MARCH5 controls lipid-induced peroxisome biogenesis. (**A**) Schematic representation of experimental protocol. (**B**) Total cell lysates from WT and MARCH5^−/−^ HeLa cells cultured for 6hr in the standard 25mM glucose-supplemented medium, or OXPHOS-stimulating glucose-deficient, 25μM palmitoyl carnitine-supplemented medium (OXPHOS) were analyzed by Western blot, as shown in the figure. Actin was a loading control. G=Glucose medium, O=OXPHOS medium, (**C**) Densitometric evaluation of the expression of representative peroxisomal proteins and components of the OXPHOS complexes in WT and MARCH5^−/−^ HeLa cells treated as in **B**. Values obtained in GLUCOSE medium were taken as 1. Mean+/−SEM. N=6-7 independent experiments. One-Sample t-test. **** p<0.0001, *** p<0.001, ** p<0.01, n.s.= no significant difference. G=Glucose medium, O=OXPHOS medium. (**D**) Number of Pex14-, PMP70- and Catalase-positive peroxisomes *per* cell in WT and MARCH5^−/−^ HeLa cells treated as in **B**. In WT cells n= 56 (PMP70), 25 (Pex14), 30 (Catalase) cells, in MARCH5^−/−^ cells n= 52 (PMP70), 25 (Pex14), and 29 (Catalase). Mean+/−SEM. One-way ANOVA. **** p<0.0001. n.s.= no significant difference. (**E**) WT (left panels) and Pex16^−/−^ (right panels) HeLa cells were fractionated into mitochondria-enriched heavy membrane (HM), light membrane (LM) and cytosolic fractions (C) followed by Western blot to detect proteins indicated in the figure. TCL: total cell lysate. Consistent data were obtained in 3 independent experiments. Red arrowheads indicate LM fractions.

Next, we tested the subcellular localization of MARCH5 in glucose and OXPHOS media-grown cells. Cell fractionations show redistribution of MARCH5 into light membrane (LM) fractions in OXPHOS medium-grown WT HeLa cells (Fig. 3E; left panels). The cell fractionation protocol we used results in the accumulation of peroxisomal proteins in a mitochondria-enriched heavy membrane (HM) and the LM fractions (Fig. 2E, 3E and Supplemental Fig. S1E). This pattern could be due to mitochondrial tethering to the peroxisomes (Chen et al., 2020) and/or the high density of some peroxisomes. Since we analyze the same amount of protein in each fraction, the peroxisomal proteins in the LM fraction could be underrepresented due to high levels of other LM organelles, such as ER and lysosomes. Nevertheless, the data show a shift of MARCH5 and peroxisome biogenesis factors Pex14 and Pex16 into LM fractions in WT cells cultured in the OXPHOS medium (Fig. 3E, left panels), suggesting that OXPHOS medium-induced peroxisome biogenesis results in an increase of LM-associated MARCH5-containing peroxisomes. We generated Pex16 knockout (Pex16^−/−^) HeLa cells to test this idea. Consistent with published works (Ott et al., 2023; Yagita et al., 2022), Pex16 knockout in HeLa cells results in a complete peroxisome deficiency (Fig S1). Notably, the shift of Pex14 and MARCH5 into LM fractions was not detected in OXPHOS medium-cultured peroxisome-deficient Pex16^−/−^ cells (Fig. 3E: right panels) indicating that MARCH5 localizes to the newly generated peroxisomes in the OXPHOS medium-treated cells and further suggesting that MARCH5 activity is required for peroxisomal biogenesis.

The potential role of MARCH5 in energy metabolism was also tested. ATP generation is the major target and outcome of metabolism regulation. To access the effect of MARCH5 deficiency on ATP generation in glucose and OXPHOS media, we measured total ATP levels (luciferase-based assay; Supplemental Fig. S2A,B), maximum ATP generation capacity of the mitochondria (luciferase-based assay; Supplemental Fig. S2C-E) (Greiser et al., 2023), and cytosolic ATP levels (time-lapse imaging of cytosolic ATP sensor cyto-iATPSnFR1.0 (Lobas et al., 2019) (Supplemental Fig. S2F-J). The cytosolic and total ATP levels were both reduced in MARCH5^−/−^ cells compared to WT HeLa cells cultured in the OXPHOS medium but not in the glucose medium (Supplemental Fig. S2). In addition, the maximum rates of mitochondrial ATP generation were ~25% lower in mitochondria isolated from MARCH5^−/−^ cells than in WT HeLa cells (Supplemental Fig. S2D). Thus, MARCH5 appears to control extramitochondrial steps in ATP generation, especially when cells are cultured in the OXPHOS medium. While currently, we cannot directly attribute this effect to MARCH5’s role in peroxisomes *per se*, MARCH5 knockout-induced deficiency in peroxisome function likely result in reduced fatty acid processing and thereby decrease in OXPHOS substrates readily available to the mitochondria. This possibility will be addressed in the future.

### MARCH5 regulates mitochondrial pre-peroxisome formation

Peroxisome biogenesis is a multi-step process comprising several peroxisome biogenesis factors. Mutations or deficiencies of several of these biogenesis factors lead to severe, but distinct peroxisomal deficiencies (complete lack of peroxisomes *versus* partial reduction and defects in maturation) (Ott et al., 2023). Until recently, it was thought that two distinct pathways form peroxisomes: the growth and fission of mature peroxisomes and *de novo* synthesis at the endoplasmic reticulum (ER) (Titorenko and Mullen, 2006). However, Sugiura *et al*. (Sugiura et al., 2017) discovered that a subset of pre-peroxisomes (immature peroxisomes that require fusion with the ER-derived pre-peroxisomes to form the metabolically active, mature organelles) originate from the mitochondria. The predominant OMM localization and function of MARCH5 (Karbowski et al., 2007; Xu et al., 2016), point to the MARCH5 function in forming the mitochondria-derived peroxisomes. We reasoned that knockouts of discrete peroxisome biogenesis factors could result in stalling peroxisome biogenesis at diverse steps. MARCH5 knockout in a background of defective peroxisomal biogenesis could overcome the potential redundancy of distinct peroxisome biogenesis factors or unmask peroxisomal targets and/or biogenesis steps requiring MARCH5 activity. To test this notion, we used Pex16^−/−^ (controls the ER-dependent steps of *de novo* peroxisome assembly (Kim et al., 2006; Wei et al., 2021)) and Pex14^−/−^ (implicated in mitochondrial steps of peroxisome biogenesis (Sugiura et al., 2017)), as well as double knockout HeLa cells, Pex16^−/−^/MARCH5^−/−^ (Pex16/M5 dko) and Pex14^−/−^/MARCH5^−/−^ (Pex14/M5 dko; Fig. 4 and Supplemental Fig.S1). Cells were analyzed by immunofluorescence and Airyscan imaging. As discussed above, no peroxisomes were detected in Pex16^−/−^ cells, and a similar phenotype was apparent in Pex16/MARCH5 dko cells (Supplemental Fig.S1B,C), indicating that MARCH5 does not affect Pex16- and ER-dependent steps in peroxisome biogenesis. On the other hand, knockout of Pex14, resulted in a marked reduction in peroxisome number, but a consistent presence of ~100 peroxisomes *per* cell was apparent (Fig. 4B,D; 922+/−175 peroxisomes *per* cell in WT, versus 103+/−33 peroxisomes in Pex14^−/−^ HeLa cells). The PMP70-positive peroxisomes in Pex14^−/−^ cells were moderately, but significantly, larger than those in WT HeLa cells (0.094+/− 0.018 μm^2^ in Pex14^−/−^ cells, *versus* 0.089+/−0.013 μm^2^ in WT cells; *p*-value=0.004; Fig. 4E). Notably, peroxisomes detected in Pex14^−/−^ cells were deficient in Catalase (Fig. 4B,F), but they invariably colocalized with the OMM marker Tom20 (Fig. 4B,J), supporting the Pex14 function in Catalase import (Okumoto et al., 2020), but also indicating that Pex14 acts in mitochondria and is required for maturation of mitochondria-derived pre-peroxisomes (Sugiura et al., 2017). Consistent with MARCH5 acting downstream of Pex14 in mitochondrial pre-peroxisome formation, the data show a complete absence of peroxisomes in Pex14/MARCH5 dko cells (Fig. 4C,D). The specificity of this phenotype and the essential role of MARCH5 E3 Ub ligase activity in mitochondrial pre-peroxisome formation is supported by the data showing that ectopic expression of WT MARCH5, but not MARCH5 mutants MARCH5^H43W^ or MARCH5^ΔCT^, rescued pre-peroxisome formation in Pex14/MARCH5 dko cells (Fig. 4G-I). Furthermore, unlike in WT and MARCH5^−/−^ cells, where only sporadic WT MYC-MARCH5 colocalization with peroxisomes was detected (Fig. 1A,D), there is a high degree of colocalization of WT MYC-MARCH5 with peroxisomes in Pex14/MARCH5 dko cells (Fig. 4G,J) indicating that Pex14 deficiency leads to stalling of WT MARCH5 on mitochondria-derived peroxisomes and suggesting molecular crosstalk between Pex14 and MARCH5. Cell fractionation further supports this possibility. As discussed above, cell fractionation of WT HeLa cells shows HM and LM localization of peroxisomal proteins (Figs. 2E, 3E, 4F,I) and deficiency of peroxisomes in Pex16^−/−^ cells eliminates peroxisome proteins in LM fraction (Fig. 3E). In line with the imaging experiments (Fig. 4A-D), reduction, or elimination of PMP70 and Pex26 was detected in LM fractions of Pex14^−/−^ and MARCH5/Pex14 dko cells, respectively (Fig. 4F). Reexpression of WT MARCH5, but not MARCH5^H43W^, nor MARCH5^ΔC^ in MARCH5/Pex14 dko cells markedly rescued PMP70 in LM fractions (Fig. 4I). On the other hand, Catalase was not affected by MARCH5 deficiency in Pex14^−/−^ (Fig. 4F) cells and reexpression in MARCH5/Pex14 dko cells (Fig. 4I), indicating that MARCH5 does not affect Pex14-dependent peroxisome maturation (e.g., Catalase import).

**Figure 4.**
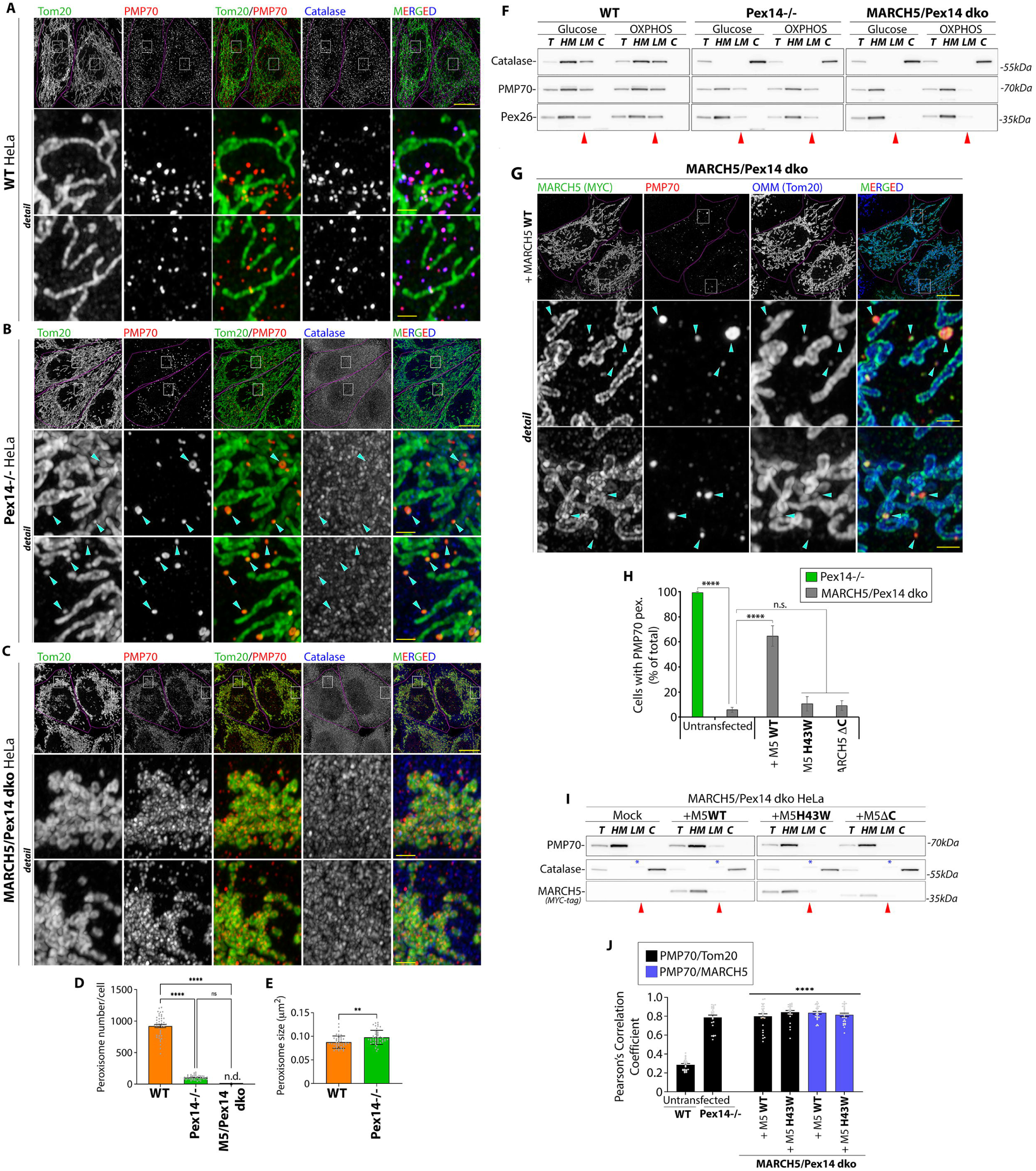
MARCH5 controls formation of mitochondria-derived pre-peroxisomes. WT (**A**) Pex14^−/−^ (**B**) and Pex14/MARCH5 dko (**C**) HeLa cells immunostained to detect Catalase (blue on merged images), PMP70 (red on merged images), and Tom20 (green on merged images) and subjected to Airy scan imaging. Bars: 10μm, and 2μm in detail images. (**D**) Number of PMP70-positive peroxisomes *per* cell in WT and Pex14^−/−^ and MARCH5/Pex14 dko HeLa cells. n=45 (pooled from 3 experiments). Mean+/− SEM. One-way ANOVA. **** p<0.0001. (**E**) Size of PMP70-positive peroxisomes in WT and Pex14^−/−^ HeLa cells. n=43 (pooled from 3 experiments). Mean+/− SEM. Two-tailed unpaired t-test. **p-value<0.01. (**F**) WT, Pex14^−/−^, and Pex14/MARCH5 dko cells were subjected to cells fractionation and Western blot analysis to detect peroxisomal proteins as indicated in the figure. Red arrowheads indicate LM membrane fractions. Consistent data were obtained in 3 out of 3 independent experiments. (**G**) MARCH5/Pex14 dko HeLa cells were transfected with MYC-WT MARCH5, immunostained to detect MARCH5 (anti-MYC; green on merged images), PMP70 (red on merged images), and OMM (Tom20; blue on merged images), followed by Airyscan imaging. Bars: 10μm, and 2μm in detail images. Arrowheads indicate PMP70-positive peroxisomes, and respective localization of MARCH5 and Tom20. (**H**) Quantification of the number of cells containing PMP70-labeled peroxisomes in Pex14^−/−^ cells and MARCH5/Pex14 dko cells, untransfected or transfected with MYC-tagged MARCH5 constructs as indicated in the figure. N=3, n=300. Mean+-SD. One-way ANOVA. **** p<0.0001. n.s.= no significant difference. (**I**) Cells fractionation of untransfected MARCH5/Pex14 dko cells, or cells transfected as indicated in the figure. *in **I** indicates a nonspecific protein. (**J**) Pearson’s colocalization coefficients representing colocalization of peroxisomes (PMP70) with the OMM (Tom20) (black bars), or peroxisomes (PMP70) with MARCH5 (blue bars) in untransfected WT and Pex14^−/−^ cells, and MARCH5/Pex14 dko cells expressing with MYC-MARCH5 and MYC-MARCH5^H43W.^ Mean+/−SEM. n=25 cells *per* condition. One-way ANOVA. **** p<0.0001.

Recent evidence indicates that *de novo* peroxisome biogenesis requires mitochondria (Schrader and Pellegrini, 2017; Sugiura et al., 2017). However, until now, the mechanism and mitochondrial factors implicated in the formation of mitochondria-dependent subsets of peroxisomes have not been determined. We report that the OMM E3 Ub ligase MARCH5 is vital for peroxisome homeostasis and biogenesis of mitochondria-derived pre-peroxisomes. Analysis of peroxisomes in two independent MARCH5 knockout cell models show a reduced number of mature peroxisomes. Supporting the idea that MARCH5 controls peroxisome biogenesis, we found that upon a shift from glycolysis-dependent growth conditions into fatty acid-supported OXPHOS-dependent growth, peroxisome biogenesis occurs in WT but not in MARCH5-deficient cells. Under these growth conditions, MARCH5 translocates to the newly formed peroxisomes. We also found that MARCH5 deficiency inhibits peroxisome formation in Pex14^−/−^ cells. Pex14 was proposed to control mitochondria-derived pre-peroxisome biogenesis (Schrader and Pellegrini, 2017; Sugiura et al., 2017). Our data support this notion and indicate that Pex14 acts upstream of MARCH5 (see Fig. 5 for the model). The mitochondria-derived pre-peroxisomes observed in Pex14^−/−^ cells were rescued in MARCH5/Pex14 dko cells by reexpression of WT MARCH5, but not MARCH5 inactive mutants, indicating that E3 Ub ligase activity of MARCH5 is critical.

**Figure 5.**
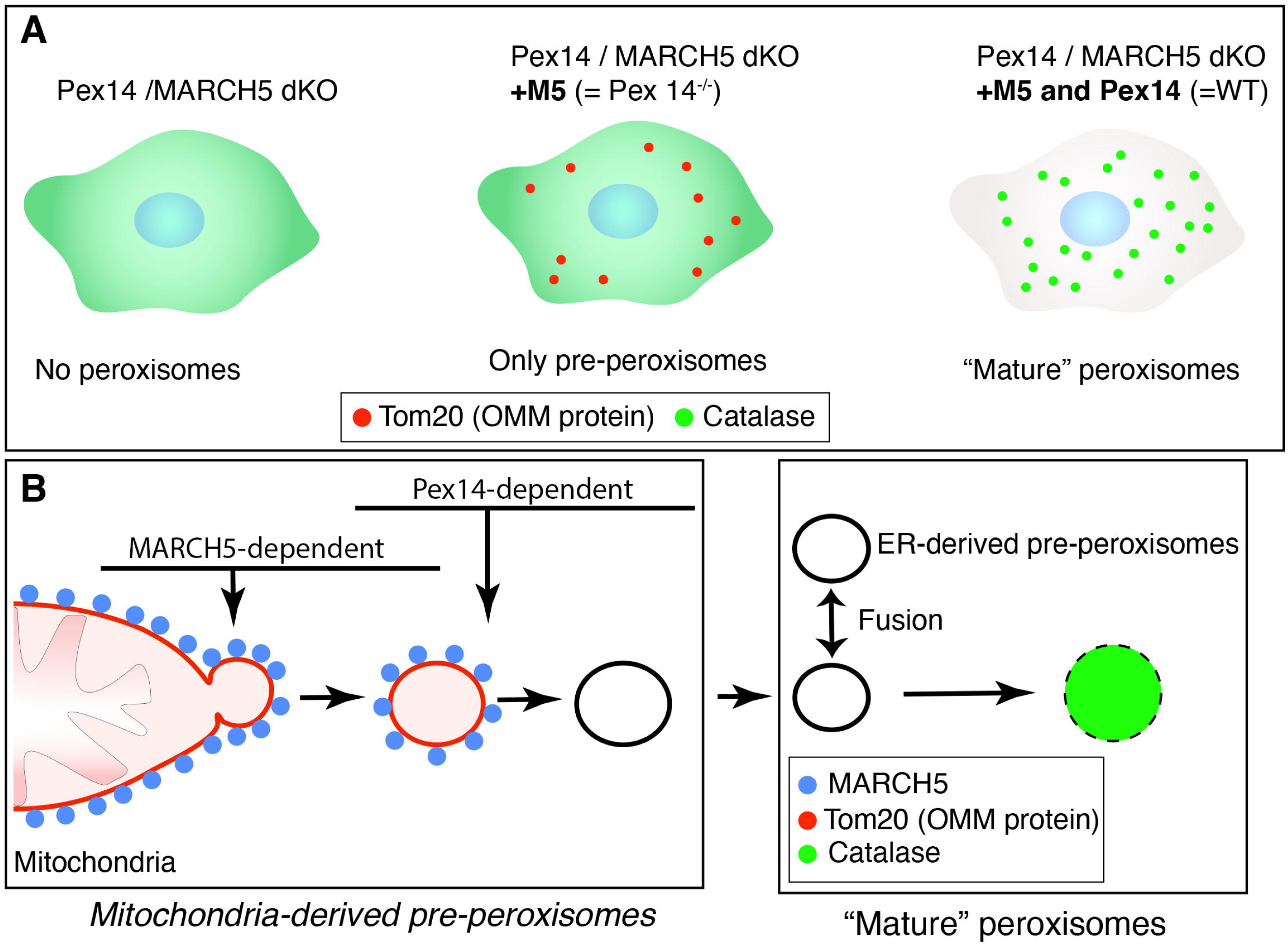
MARCH5 controls formation of mitochondria-derived pre-peroxisomes (MODEL) (**A**) Double knockout of Pex14 and MARCH5 result in complete elimination of peroxisomes, while single Pex14 knockout show mitochondria-derived pre-peroxisomes but not mature peroxisomes, indicating that MARCH5 controls early steps of mitochondrial pre-peroxisome biogenesis, upstream of Pex14 (**B**).

## Material and Methods

### Cells and cell culture

HeLa and HCT116 cells were maintained in Dulbecco Modified Eagle Medium (DMEM; Invitrogen) supplemented with 10% fetal bovine serum (FBS; Sigma), non-essential amino acids (Sigma), sodium pyruvate (Invitrogen), and penicillin/streptomycin (Invitrogen). Cells were maintained in 5% CO_2_ at 37°C.

### Knockout cells, DNA constructs, and transfections

MARCH5^−/−^ HCT116 and Drp1^−/−^ HeLa cells were reported previously (Cherok et al., 2017; Oshima et al., 2021; Xu et al., 2016). MARCH5^−/−^, Pex14^−/−^, Pex16^−/−^, Pex26^−/−^, Pex14/MARCH5 dko, Pex16/MARCH5 dko and Pex26/MARCH5 dko HeLa cells were generated using double nickase plasmids from Santa Cruz Biotechnology. Cells grown on 6-well plates were transfected with either single or co-transfected with two plasmids (ratio 1:1). At 48hr after transfection, cells were treated with fresh media containing 3μg/ml puromycin and grown in puromycin-containing media for 4 days, with daily media change. Then, cells were split into 25-cm cell culture plates and cultured in puromycin-deficient media until colonies were formed, followed by the additional 72-hr culture in puromycin-supplemented media. This treatment eliminated puromycin-sensitive cells carried over after the first puromycin selection, which usually represented ~70% of the colonies. Puromycin-resistant colonies were sequentially transferred into 24-well plates and then into duplicate 6-well plates. Cells from one 6-well plate were used for knockout verifications (Cherok et al., 2017; Xu et al., 2016), and from another for propagation. Mammalian expression vectors encoding MYC-tagged dominant negative mutant of Drp1 (MYC-Drp1^K38A^), MYC-tagged WT MARCH5, MARCH5^H43W^, MARCH5^ΔCT^ were reported (Cherok et al., 2017; Oshima et al., 2021; Xu et al., 2016). The cytosolic ATP sensor cyto-iATPSnFR1.0 was a gift from Baljit Khakh (Addgene plasmid# 102550; http://n2t.net/addgene:102550; RRID:Addgene_102550) (Lobas et al., 2019). Cells were transfected with Lipofectamine 3000 (Thermo Fisher Scientific) according to the manufacturer’s instructions. Cells were used at ~12-20 h after transfection.

### Cell lysates, cell fractionation, and Western blot

For total-cell lysates, cells were collected by scraping into ice-cold PBS, washed, and suspended in ice-cold PBS. Cell suspensions (100-200μl) were lysed in the same volumes of 2 x SDS sample buffer (Thermo Fisher Scientific) supplemented with 5% β-mercaptoethanol (Millipore) and incubated at 100°C for 10 min, as described (Cherok et al., 2017; Xu et al., 2016). Cell fractionation was performed as previously described (Cherok et al., 2017; Oshima et al., 2021). Cells were washed once with ice-cold PBS and scraped into 15-ml tubes in ice-cold PBS; this was followed by centrifugation at 500 × g for 5 min. The cell pellets were resuspended in ~3 volumes of fractionation buffer (10 mM HEPES, 10 mM NaCl, 1.5 mM MgCl_2_, 5mM N-ethylmaleimide (NEM), and protease inhibitors (Sigma). Cells were then passed 15 times through a 25-G needle attached to a 1-ml syringe to disrupt cell membranes. This suspension was centrifuged at 2500 × g at 4°C for 5 min to remove unbroken cells and cell debris. The supernatant was centrifuged at 6000 × g at 4°C for 10 min to pellet the heavy membrane (HM) fraction. The supernatants were centrifuged at 21,000 × g at 4°C for 10 min to pellet the light membrane (LM) fraction. To reduce cytosolic and HM contaminations, the HM and LM fractions were washed in ice-cold PBS supplemented with protease inhibitors (Sigma) and recentrifuged at 8000 × g, and 21,000 × g, respectively. Protein concentrations were measured directly in the samples using a NanoDrop 1000 spectrophotometer (Thermo Fisher Scientific). 50μg of protein per sample was separated on 4-20% or 10-20% gradient Novex Tris-glycine polyacrylamide gels (Thermo Fisher Scientific) transferred onto polyvinylidene fluoride membranes (Bio-Rad Laboratories). Membranes were blocked in 5% blocking-grade nonfat dry milk (Bio-Rad Laboratories) in PBS-Tween20 and incubated with primary antibodies overnight at 4°C, followed by horseradish peroxidase-conjugated anti-mouse (Cell Signaling) or anti-rabbit (Cell Signaling) secondary antibodies for 60 min at RT. Blots were developed with Super Signal West Pico ECL (Thermo Fisher Scientific). Blots were imaged using Amersham Imager 600 chemiluminescence imager (GE Healthcare Life Sciences). Primary antibodies used for Western blotting and their dilutions are listed in the Reagents and Resources table. Densitometric evaluations of protein expression were performed using the ImageJ64 image analysis software (NIH, Bethesda, MD), as reported (Cherok et al., 2017; Oshima et al., 2021).

**Table 1.**
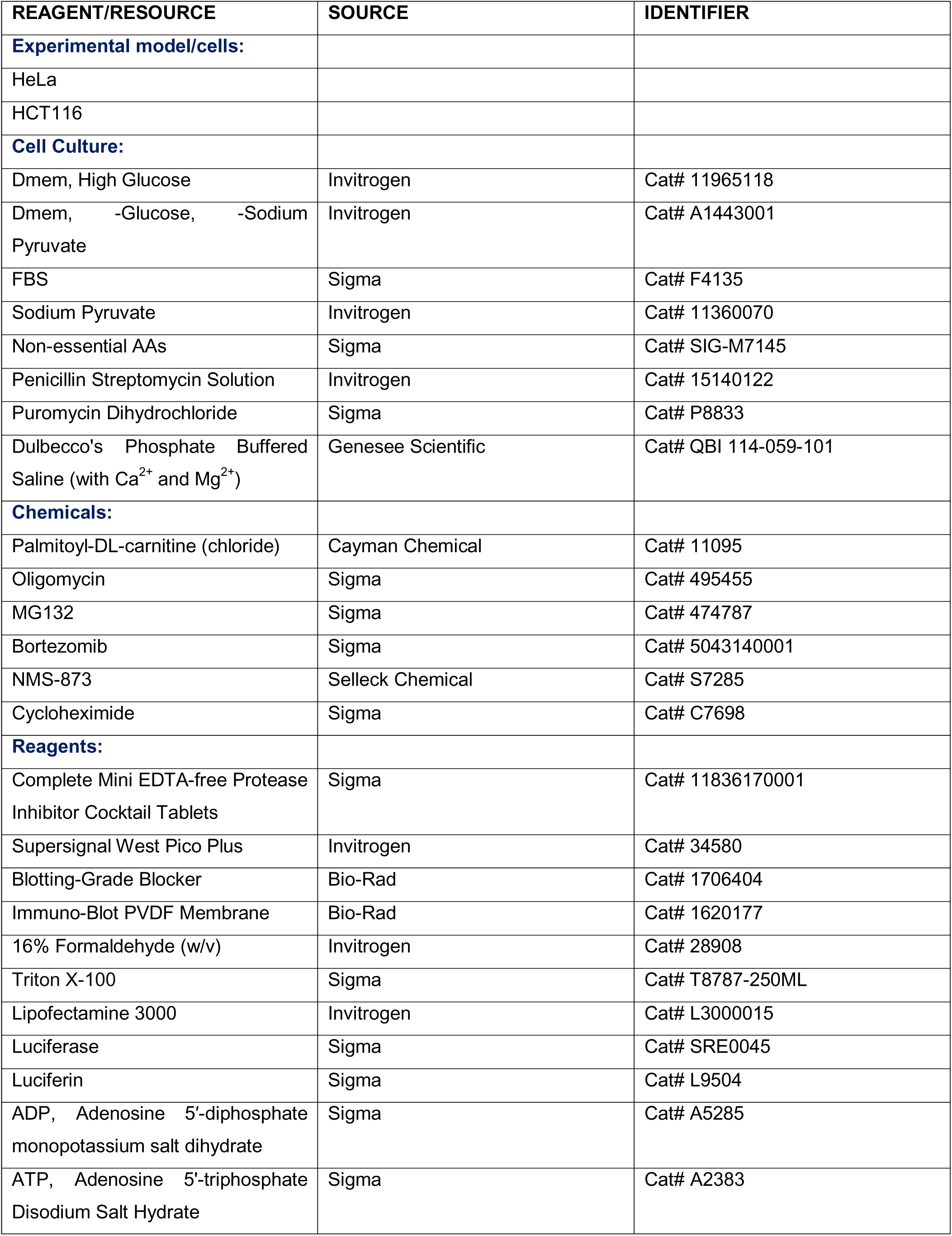
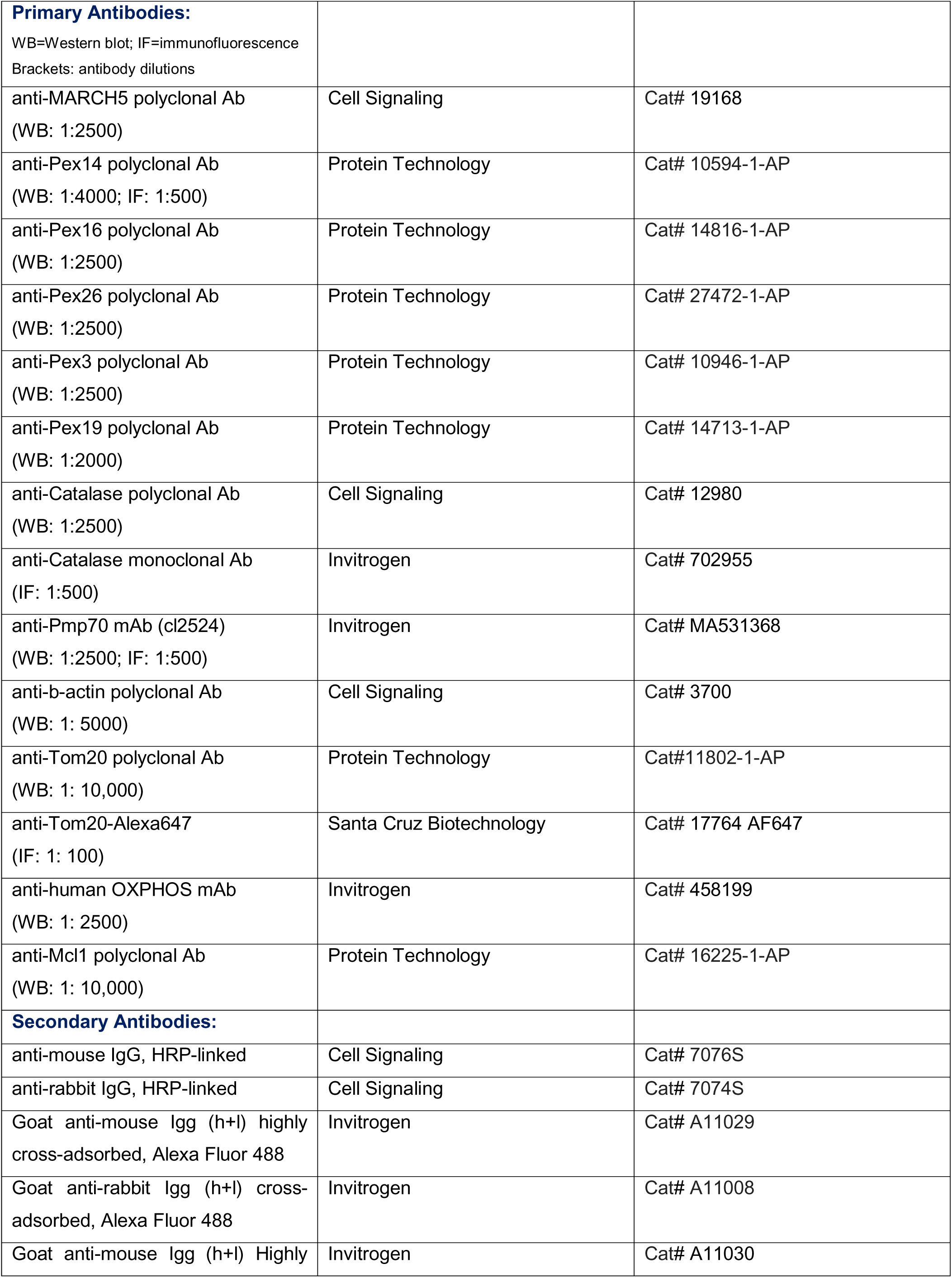
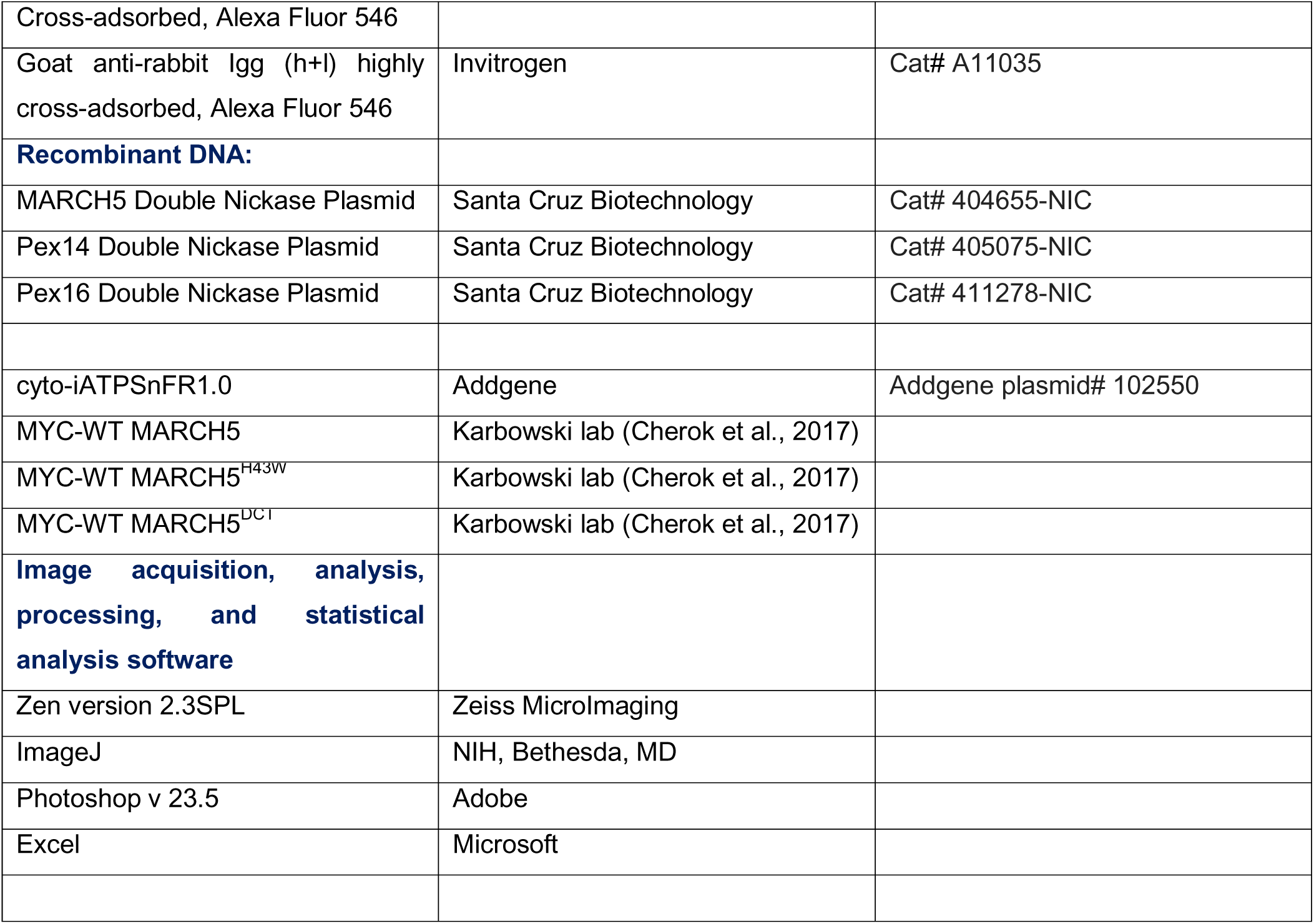
Reagents and resources used in the study.

| REAGENT/RESOURCE | SOURCE | IDENTIFIER |
| --- | --- | --- |
| **Experimental model/cells:** |  |  |
| HeLa |  |  |
| HCT116 |  |  |
| **Cell Culture:** |  |  |
| Dmem, High Glucose | Invitrogen | Cat# 11965118 |
| Dmem, -Glucose, -Sodium Pyruvate | Invitrogen | Cat# A1443001 |
| FBS | Sigma | Cat# F4135 |
| Sodium Pyruvate | Invitrogen | Cat# 11360070 |
| Non-essential AAs | Sigma | Cat# SIG-M7145 |
| Penicillin Streptomycin Solution | Invitrogen | Cat# 15140122 |
| Puromycin Dihydrochloride | Sigma | Cat# P8833 |
| Dulbecco's Phosphate Buffered Saline (with Ca<sup>2+</sup> and Mg<sup>2+</sup>) | Genesee Scientific | Cat# QBI 114-059-101 |
| **Chemicals:** |  |  |
| Palmitoyl-DL-carnitine (chloride) | Cayman Chemical | Cat# 11095 |
| Oligomycin | Sigma | Cat# 495455 |
| MG132 | Sigma | Cat# 474787 |
| Bortezomib | Sigma | Cat# 5043140001 |
| NMS-873 | Selleck Chemical | Cat# S7285 |
| Cycloheximide | Sigma | Cat# C7698 |
| **Reagents:** |  |  |
| Complete Mini EDTA-free Protease Inhibitor Cocktail Tablets | Sigma | Cat# 11836170001 |
| Supersignal West Pico Plus | Invitrogen | Cat# 34580 |
| Blotting-Grade Blocker | Bio-Rad | Cat# 1706404 |
| Immuno-Blot PVDF Membrane | Bio-Rad | Cat# 1620177 |
| 16% Formaldehyde (w/v) | Invitrogen | Cat# 28908 |
| Triton X-100 | Sigma | Cat# T8787-250ML |
| Lipofectamine 3000 | Invitrogen | Cat# L3000015 |
| Luciferase | Sigma | Cat# SRE0045 |
| Luciferin | Sigma | Cat# L9504 |
| ADP, Adenosine 5'-diphosphate monopotassium salt dihydrate | Sigma | Cat# A5285 |
| ATP, Adenosine 5'-triphosphate Disodium Salt Hydrate | Sigma | Cat# A2383 |
| **Primary Antibodies:**<br>WB=Western blot; IF=immunofluorescence<br>Brackets: antibody dilutions |  |  |
| anti-MARCH5 polyclonal Ab<br>(WB: 1:2500) | Cell Signaling | Cat# 19168 |
| anti-Pex14 polyclonal Ab<br>(WB: 1:4000; IF: 1:500) | Protein Technology | Cat# 10594-1-AP |
| anti-Pex16 polyclonal Ab<br>(WB: 1:2500) | Protein Technology | Cat# 14816-1-AP |
| anti-Pex26 polyclonal Ab<br>(WB: 1:2500) | Protein Technology | Cat# 27472-1-AP |
| anti-Pex3 polyclonal Ab<br>(WB: 1:2500) | Protein Technology | Cat# 10946-1-AP |
| anti-Pex19 polyclonal Ab<br>(WB: 1:2000) | Protein Technology | Cat# 14713-1-AP |
| anti-Catalase polyclonal Ab<br>(WB: 1:2500) | Cell Signaling | Cat# 12980 |
| anti-Catalase monoclonal Ab<br>(IF: 1:500) | Invitrogen | Cat# 702955 |
| anti-Pmp70 mAb (cl2524)<br>(WB: 1:2500; IF: 1:500) | Invitrogen | Cat# MA531368 |
| anti-b-actin polyclonal Ab<br>(WB: 1: 5000) | Cell Signaling | Cat# 3700 |
| anti-Tom20 polyclonal Ab<br>(WB: 1: 10,000) | Protein Technology | Cat#11802-1-AP |
| anti-Tom20-Alexa647<br>(IF: 1: 100) | Santa Cruz Biotechnology | Cat# 17764 AF647 |
| anti-human OXPHOS mAb<br>(WB: 1: 2500) | Invitrogen | Cat# 458199 |
| anti-Mcl1 polyclonal Ab<br>(WB: 1: 10,000) | Protein Technology | Cat# 16225-1-AP |
| **Secondary Antibodies:** |  |  |
| anti-mouse IgG, HRP-linked | Cell Signaling | Cat# 7076S |
| anti-rabbit IgG, HRP-linked | Cell Signaling | Cat# 7074S |
| Goat anti-mouse IgG (h+l) highly cross-adsorbed, Alexa Fluor 488 | Invitrogen | Cat# A11029 |
| Goat anti-rabbit IgG (h+l) cross-adsorbed, Alexa Fluor 488 | Invitrogen | Cat# A11008 |
| Goat anti-mouse IgG (h+l) Highly | Invitrogen | Cat# A11030 |
| Cross-adsorbed, Alexa Fluor 546 |  |  |
| Goat anti-rabbit IgG (h+l) highly cross-adsorbed, Alexa Fluor 546 | Invitrogen | Cat# A11035 |
| **Recombinant DNA:** |  |  |
| MARCH5 Double Nickase Plasmid | Santa Cruz Biotechnology | Cat# 404655-NIC |
| Pex14 Double Nickase Plasmid | Santa Cruz Biotechnology | Cat# 405075-NIC |
| Pex16 Double Nickase Plasmid | Santa Cruz Biotechnology | Cat# 411278-NIC |
| cyto-iATPSnFR1.0 | Addgene | Addgene plasmid# 102550 |
| MYC-WT MARCH5 | Karbowski lab (Cherok et al., 2017) |  |
| MYC-WT MARCH5<sup>H43W</sup> | Karbowski lab (Cherok et al., 2017) |  |
| MYC-WT MARCH5<sup>DC1</sup> | Karbowski lab (Cherok et al., 2017) |  |
| **Image acquisition, analysis, processing, and statistical analysis software** |  |  |
| Zen version 2.3SPL | Zeiss MicroImaging |  |
| ImageJ | NIH, Bethesda, MD |  |
| Photoshop v 23.5 | Adobe |  |
| Excel | Microsoft |  |

### Immunofluorescence

Immunofluorescence was performed as previously described (Cherok et al., 2017; Xu et al., 2011). Briefly, cells grown in 2- or 4-well chamber slides (model 1 German borosilicate; Lab-Tek; VWR) were fixed with freshly prepared 4% formaldehyde in PBS solution (using 16% Methanol-free Formaldehyde; Thermo Fisher Scientific) for 20 min at RT, then permeabilized with permeabilization buffer (PB; 0.15% Triton X-100 in PBS) for 20 min at RT. After blocking with blocking buffer (BB; PB supplemented with 7.5% BSA) for 45 min, samples were incubated with primary antibodies suspended in BB for 90 min at RT, followed by 3 washes with BB and incubation with secondary antibodies diluted in BB for 60 min at RT. Primary antibodies and their dilutions are listed in the Reagents and Resources table. Secondary antibodies were highly cross-absorbed goat anti-mouse Alexa Fluor-488 and goat anti-rabbit Alexa Fluor-546 (1:1000; both from Thermo Fisher Scientific). In triple labeling experiments, cells were immunostained, as above, followed by 3 washes with BB and incubation with anti-Tom20 antibodies conjugated with Alexa 647 fluorophore (Santa Cruz Biotechnology Inc.; 1:100) for 90 min at RT. After 3 washes with PBS, cells were subjected to Airyscan imaging. Immunolabeled cells on 4-well chamber slides were stored in PBS at 4°C and imaged within 10 days after processing.

### Image acquisition, analysis and processing

Images were acquired with a Zeiss LSM 880 microscope (Zeiss MicroImaging) equipped with an Airyscan superresolution imaging module, using 63/1.40 Plan-Apochromat Oil DIC M27 objective lens (Zeiss MicroImaging), as described (Cherok et al., 2017; Oshima et al., 2021). The 488-nm Argon laser line, 561-nm DPSS 561 laser, and 633-nm HeNe 633 laser were used to detect Alexa-488, Alexa-546, and Alexa-647, respectively. The *z*-stacks covering the entire depth of cells with intervals of 0.018μm were acquired, followed by Airyscan image processing (set at 6) and generation of maximum intensity projection images. The lateral resolution of the resulting images was ~120nm.

Image processing and analyses were done using ZEN software (version 2.3SPL, Zeiss MicroImaging) and ImageJ (NIH, Bethesda, MD) image acquisition and processing software. Colocalization/overlap, indicated by Pearson’s correlation coefficient (R), was determined from maximum intensity projections of the Airyscan processed images using a “colocalization” function of the ZEN software. For quantifications of fluorescence profiles, single z-section Airyscan-processed images were used. The profiles along peroxisomes were generated from the 2-channel images using the “profile” option available in the ZEN software. For peroxisome number and size, single cells in maximum-intensity projection Airyscan images were outlined, cropped, and converted into binary images using ImageJ, and analyzed using the “particle analysis” module of the software. The data were tabularized and transferred to Microsoft Excel software (Microsoft) for further analyses. Image cropping and global brightness and contrast adjustments were performed for presentation using Adobe Photoshop CS6 software (Adobe Systems).

### Time-lapse live cell imaging of cytosolic ATP

For time-lapse imaging, cells were grown on 4-well chamber slides (model 1 German borosilicate; Lab-Tek; VWR) to 30-50% confluency. Cells were transfected with cytosolic ATP sensor cyto-iATPSnFR1.0 <14 hr before analyses. In all time-lapse experiments chamber slides were mounted on the environmental control chamber (Stagetop TIZW Series, Neco Incubation System with sensor feedback system) attached to Zeiss LSM 880 microscope (Zeiss MicroImaging) set at 5% CO_2_ and 37°C and imaged in Phenol Red-free DMEM, 10% fetal bovine serum (FBS; Sigma), non-essential amino acids (Sigma), penicillin/streptomycin (Invitrogen), supplemented with 0-25mM glucose, galactose, or 25μm Palmitoyl-DL-carnitine, as described in the text.

### Total and mitochondrial ATP measurements

For total ATP measurements, cells were grown overnight in 10-cm cell culture plates in 25mM glucose medium to 80% confluence. Cells were washed and incubated in the starvation medium (DPBS) for 60min to reduce cellular ATP (Zeidler et al., 2017) followed by growth for 6hr in 5% CO_2_ at 37°C in DPBS anew, 4.5mM galactose medium, 25mM glucose medium, and 4.5mM galactose media supplemented with 25µM pLcar, and 25mM glucose supplemented with 25µM pLcar. To inhibit mitochondrial ATP generation, each condition (except the DPBS condition) was also subjected to treatment with 5µM oligomycin (Sigma). ATP extractions were performed using a protocol described by Yang *et al*. (Yang et al., 2002), with modification. Cells were collected, transferred to 2ml Eppendorf tubes, and exposed to boiling deionized water containing 0.5% Triton-X 100 (Sigma). Denatured protein clumps were broken down by 10 passages through a 25 G x 5/8 needle attached to 1 ml syringe. Samples were centrifuged at 20,000 x g for 10 min, at 4°C. The supernatants were then transferred to a new tube and used in measurements. Pellets were solubilized in 2 x SDS PAGE sample buffer supplemented with 5% β-mercaptoethanol, incubated at 100°C for 10 min, and centrifuged at 20,000 x g for 5 minutes at 4°C. The supernatants were transferred to new tubes and used for protein measurements. Mitochondrial ATP measurements were performed as previously described (Shimada et al., 2022; Wescott et al., 2019). Cells were grown to confluency on 15-cm plates, the growth media was discarded, and the plates were washed twice with ice-cold PBS supplemented with 1 mM EGTA (pH 7.4). The cells were then harvested using a cell scraper, transferred to 50 mL conical tubes, and centrifuged at 500 x g to pellet the cells and discard the PBS solution. Cells were then resuspended in ice-cold isolation buffer (IB) containing (in mM): KCl 100, MOPS 50, MgSO_4_ 5, EGTA 2, K_2_HPO_4_ 10. The cell suspension was washed twice with IB by centrifugation at 500 x g to pellet the cells and discard the supernatant. The remainder of the preparation was conducted in a cold room (4°C). 10 ml of IB containing cells underwent 15 repetitive homogenizations with a 1-μm clearance pestle at low speed. The homogenate was centrifuged for 8 min at 600 x g at 4°C, after which the supernatant was transferred to a new centrifuge tube. These homogenizations of the cell pellet and centrifugation to pellet cells were repeated 3 more times. The supernatant from four homogenizations and centrifugation was pooled together, transferred to new centrifuge tubes, and spun at 600 x g for 8 min at 4°C to pellet cell debris The supernatant was transferred to a clean centrifuge tube and spun at 600 x g again to pellet cell debris for 8 min. The supernatant was transferred to a clean centrifuge tube and spun at 3200 x g to pellet mitochondria for 12 min. The supernatant was discarded, and the pellet (the mitochondria sample) was resuspended in IB warmed to 30°C and supplemented with Na-Pyruvate (10 mM). After 10 min incubation at room temperature, the suspension was spun at 3200 x g for 12 min to pellet the mitochondria. The supernatant was discarded, and the pellet (the mitochondria sample) was resuspended in ice-cold IB supplemented with Na-Pyruvate (10 mM) and kept on ice for 30 min and centrifuged at 3200 x g for 12 min to pellet mitochondria. The pellet was resuspended in ice cold resuspension buffer (RB1) base solution containing (in mM): KCl 100, MOPS 50, K_2_HPO_4_ 1, supplemented with Na-Pyruvate (10 mM), EGTA (40 μM). The mitochondria were centrifuged at 3200 x g at 4°C for 12 min and resuspended in RB2, which is RB supplemented with Na-Pyruvate (1 mM) and EGTA (40 μM). The mitochondria were centrifuged at 3200 x g for 12 min, and a final resuspension and pelleting were done using RB3, consisting of RB and EGTA (40 μM). The concentration of mitochondria was quantified by Lowry assay. The high purity of mitochondria isolated via this procedure was previously shown (Shimada et al., 2022; Wescott et al., 2019). Mitochondria were used within 4 hours of isolation. Measurements of total cellular ATP and mitochondrial ATP production rate were carried out using a BMG LABTECH CLARIOstar plate reader. Mitochondria (0.1 mg per mL) were mixed in ATP production assay buffer (AB) consisting of (in mM): K-Gluconate 130, KCl 5, K_2_HPO_4_ 1 or 10, MgCl_2_ 1, HEPES 20, EGTA 0.04, BSA 0.5 mg/mL, D-Luciferin (Sigma) 0.005, Luciferase (Sigma) 0.001 mg/mL, pH 7.2. A luminescence standard curve was performed daily over a range of 100nM to 1 mM ATP with Oligomycin A (15 μM) treated mitochondria. The mitochondria were supplemented for 2 minutes prior to the start of the assay in 1 mM Pyruvate and 0.5 mM Malate, 1 mM Glutamate and 0.5 mM Malate, or 0.1 mM palmitoylcarnitine + 2.5 mM malate. Assays were initiated by injection of 20 μL ADP (5.0 mM) and 80 μL luciferin/luciferase in AB to bring the final volume to 200 μL. Luminescence signal was recorded for 20 seconds with 1 second integration. ATP production rates were scaled to nMol ATP per sec per mg mitochondrial protein (nMol S^−1^ mg^−1^).

### Statistical calculations

Unless otherwise indicated, data was analyzed using unpaired, two-tail Student’s test; for multiple-group comparisons, data were analyzed using one-way ANOVA with Bonferroni correction (α=0.05) to access significance. All error bars are expressed as SEM. P-values < 0.05 were considered significant. “N” indicates independent experiments; “n” indicates repeats within each experiment.

## Abbreviations

OMM: outer mitochondrial membrane
Ub: ubiquitin
Pex: peroxin
pLcar: palmitoyl-L-carnitine
OXPHOS: oxidative phosphorylation

## Acknowledgements

We would like to thank Dr. W. Jonathan Lederer for providing ample access to state-of-the-art imaging systems and analytic resources. Research reported in this publication was supported by the National Institutes of Health under Award Numbers R01GM129584 (MK) and by the Center for Biomedical Engineering and Technology (BioMET), University of Maryland, Baltimore.

## Author contribution

NV, YO, EC, LB, and MK conducted the experiments and analysed data, AN generated some DNA constructs, MK designed the study, supervised the project, and wrote the manuscript. All authors commented on and critically influenced the current version of the manuscript.

