## Supplementary figures and images for "Outer mitochondrial membrane E3 Ub ligase MARCH5 controls mitochondrial steps in peroxisome biogenesis"

### Supplemental Figure 1

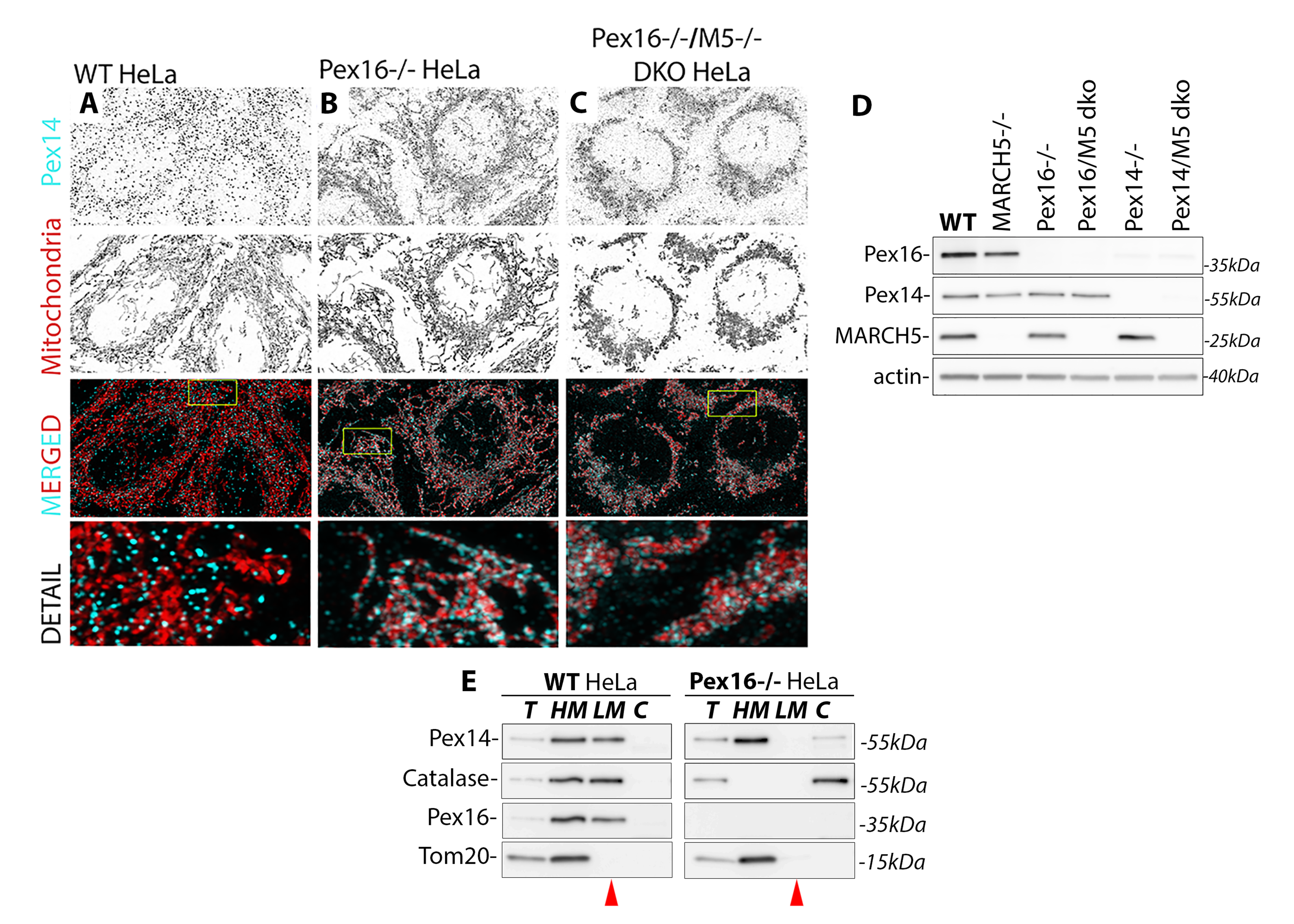

### Supplemental Figure 2

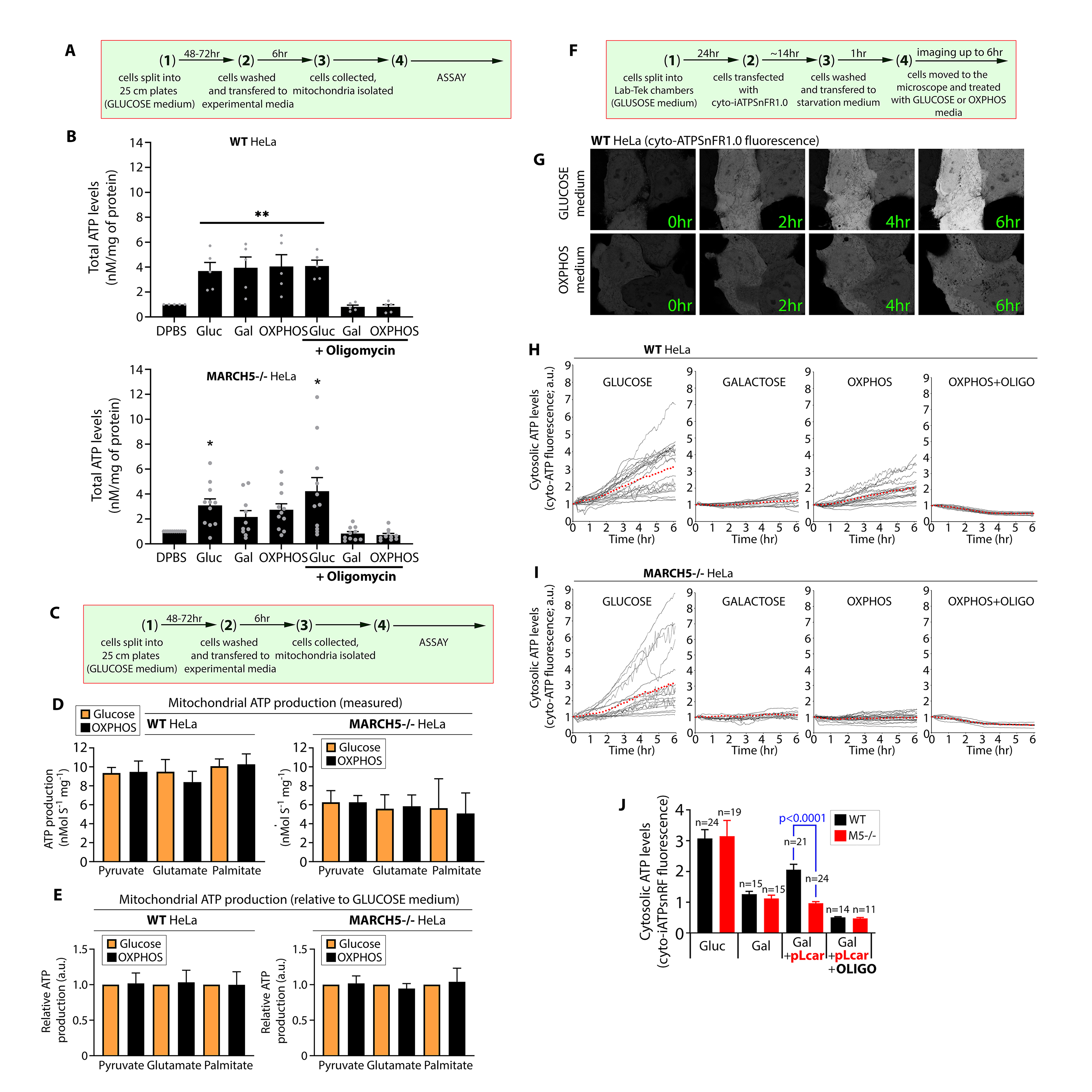
