## Supplemental Text for "Outer mitochondrial membrane E3 Ub ligase MARCH5 controls mitochondrial steps in peroxisome biogenesis"

**Supplemental Figure legends**

**Figure S1. Peroxisomes in** **Pex16 and Pex16/MARCH5 knockout HeLa cells.**

Typical images of WT (**A**), Pex16^-/-^ (**B**) and Pex16/MARCH5 dko (**C**) HeLa cells that were immunostained to detect Pex14 (light blue on merged images) and Tom20 (red on merged images). (**D**) Total cell lysates of WT, MARCH5^-/-^, Pex16^-/-^, Pex16/MARCH5 dko, Pex14^-/-^, and Pex14/MARCH5 dko were analyzed by Western blot as shown in the figure. Actin is a loading control. (**E**) WT and Pex16^-/-^ HeLa cells were fractionated into mitochondria-enriched heavy membrane (HM), light membrane (LM) and cytosolic fractions (C) followed by Western blot to detect proteins indicated in the figure. Red arrows indicate shift in Pex14, Catalase, and Pex16 protein between WT and Pex16^-/-^ in LM fraction. T: total cell lysate.

**Figure S2. Total, mitochondrial, and cytosolic ATP generation/levels in glucose and OXPHOS media cultured WT and MARCH5^-/-^ HeLa cells.**

Total (**A**,**B**), mitochondrial (**C**-**E**), and cytosolic (**F**-**J**) ATP a in WT and MARCH5^-/-^ cells cultured in glucose and OXPHOS media, as indicated in the figure, were tested. Experimental setups for each subset of ATP are shown in **A**, **C**, and **F** (light green rectangles). (**B**)Total ATP was measured using bioluminescence assay. N=5 (WT cells) and 11 (MARCH5^-/-^ cells). One-way ANOVA. ** p<0.01, *p<0.05.

(**C**-**E**) Maximal mitochondrial ATP production capacity in WT and MARCH5^-/-^ HeLa cells. Maximal mitochondrial ATP production was stimulated by 0.5 mM ADP and different metabolic substrates, as indicated in the figure, and measured by luciferase activity. In **D**, the measured ATP generation rates calculated as (nMoles of ATP x S^-1^ x mg protein^-1^) in WT (left panels) and MARCH5^-/-^ (right panels) HeLa cells are shown. In **E**, mitochondrial ATP generation in OXPHOS medium-pretreated cells relative to the values obtained in mitochondria from Glucose medium-pretreated mitochondria (taken as 1) is shown. Mean±SEM. N=8.

(**F**-**J**) Cytosolic ATP levels were accessed using time-lapse imaging of cells expressing cytosolic ATP sensor cyto-iATPSnFR1.0 (Lobas et al., 2019). At <24hr after transfection, to reduce ATP levels, cells were starved by incubation in DPBS (with Ca^2+^ and Mg^2+^; starvation medium) for 60min, followed by transfer into glucose or OXPHOS media and time-lapse image acquisition. (**G**) Typical images of WT HeLa cells expressing cyto-iATPSnFR1.0 in glucose (top panels) and OXPHOS medium (bottom panels) at 0, 2, 4, and 6hr of imaging. Changes in the cyto-iATPSnFR1.0 fluorescence in WT HeLa (**H**), and MARCH5^-/-^ (**I**) HeLa cells treated as indicated in the figure are plotted as a function of time. n-indicates the number of analyzed cells *per* each condition. Black lines: single-cell traces, red lines: mean of all single cell experiments. (**J**) Quantification of cyto-iATPSnFR1.0 fluorescence at 6hr of treatment, as indicated. The value at time 0hr was taken as 1. Mean±SEM. N indicated in the figure. One-way ANOVA.
